# Sentence processing is modulated by the current linguistic environment and a priori information: An fMRI study

**DOI:** 10.1101/416404

**Authors:** K. Weber, C. Micheli, E. Ruigendijk, J.W. Rieger

## Abstract

Words are not processed in isolation but in rich contexts that are used to modulate and facilitate language comprehension. Here, we investigate distinct neural networks underlying two types of contexts. Firstly, the current linguistic environment, presented as the relative frequencies of two syntactic structures (prepositional object (PO) and double-object (DO)), which would either follow everyday linguistic experience or not. Secondly, preference towards one or the other structure depending on the verb; learned in everyday language use and stored in memory. German participants were reading PO and DO sentences in German while brain activity was measured with functional magnetic resonance imaging. Firstly, the anterior cingulate cortex (ACC) showed a pattern of activation that integrated the current linguistic environment with everyday linguistic experience. When the input did not match everyday experience, the unexpectedly frequent structure showed higher activation in the ACC than the other conditions and more connectivity from the ACC to posterior parts of the language network. Secondly, verb-based surprisal of seeing a structure given a verb (PO verb preference but DO structure presentation) resulted, within the language network (left inferior frontal and left middle/superior temporal gyrus) and the precuneus, in increased activation compared to a predictable situation. In conclusion, 1) beyond the canonical language network, brain areas engaged in cognitive control, such as the ACC, might use the statistics of syntactic structures to facilitate language comprehension, 2) the language network is directly engaged in processing verb preferences. These two networks show distinct influences on sentence processing.

## Introduction

When we process language, when we try to extract meaning from texts, during conversation, in any situation in which we work with language, we use many different sources of information, from the preceding words to speaker identity, to make language processing fast and efficient (Christiansen & Chater, 2016; Kuperberg & Jaeger, 2016; Pickering & Garrod, 2007). We adapt to the statistics of the current or recent environment (Fine, Jaeger, Farmer, & Qian, 2013; Segaert, Weber, Cladder-Micus, & Hagoort, 2014; Wells, Christiansen, Race, Acheson, & MacDonald, 2009) as well as using information stored in memory about the general frequency of occurrence of words, structures and their co-occurrence. The adaptation to these two types of information occurs on different time scales: a short time scale of the experimental context with different relative frequencies of sentence structures and a long time scale of verb biases learned over a lifetime of language use. In the present study we investigated how the brain networks involved in processing the preceding context and stored frequency information modulate language processing and how they might interact. This study will thus investigate the invariance and variability of the language network (and beyond) in processing different types of contextual and predictive information.

The brain adapts to the statistics of the input, including the frequencies of semantic or syntactic features. As Neely (1991) already showed a few decades ago, semantic priming effects are affected by the context, i.e. semantic priming effects are larger if they occur in contexts with a lot of semantically related pairs. Also syntactic priming effects are influenced by changes in the statistics of the input (Segaert, Menenti, Weber, & Hagoort, 2011), more specifically, exposure to a large number of sentences of one particular structure will modulate the magnitude of the syntactic priming effects for that structure (decrease in magnitude) as well as its infrequent counterpart (increase in magnitude). Thus, the brain is sensitive to the proportion of different linguistic features such as words, semantic relations and syntactic structures in the input and can use this information to modulate language processing. These changes in the overall input statistics, e.g. an increased likelihood of occurrence of a certain syntactic structure, lead to predictions of encountering more of these structures and can be used to facilitate processing.

Next to adaptation to syntactic structure we also generate predictions based on prior experience with the language that we have stored in memory. We have learned that certain sentence structures are used more frequently but also that certain words, such as verbs carry different likelihoods of being paired with certain syntactic structures. Prepositional object (PO) structures, such as ‘The girl gave the flower to the boy’ and double object (DO) structures such as ‘The girl gave the boy the flower’ are ditransitive sentences that form a syntactic alternation, they carry the same meaning but are expressed with two different grammatical structures. Different verbs have different preferences for one or the other structure (see Table 1 for examples), and we gain this knowledge during our experience with the language. It has been shown that these verb-biases towards syntactic structure modulate sentence processing: for example predictive effects based on verb-based preferences were shown in a visual world paradigm (Arai & Keller, 2013), verb-biases influence ambiguity resolution (Garnsey, Pearlmutter, Myers, & Lotocky, 1997) and verb-biases modulate syntactic priming effects (Bernolet & Hartsuiker, 2010; Melinger & Dobel, 2005; Segaert et al., 2014). Therefore, this information about the frequency of co-occurrence of verb and syntactic structure must be stored in memory and can thus be used to predict which syntactic structure is likely to come up next. Moreover, different languages adopt different statistics with regard to the general use of one structure over the other. For example in the language tested in the present experiment, German, the double-object construction is overall preferred over the prepositional object construction.

**Table 1.** Verbs and nouns used in the experimental sentences and verb syntactic preference values. In combination with the potential nouns for the agent and recipient (man, woman, boy, girl) these formed the different sentences. The values in the third column were used to calculate verb based syntactic surprisal effects for the fMRI regressor (see results section). Note that the values for the verb preference differ (different preference) between the pre- and the post-test (column 2 and 3) in a few of the cases. For the analysis we used column 3, which reflects the verb preferences of the group of participants we tested.

| Verb (E translation) | DO-preference previous study (how normal?) | DO-preference post-test (sentence continuation) | Preference based on pretest | Preference based on posttest | preposition | themes (E translation) |
| --- | --- | --- | --- | --- | --- | --- |
| bereiten (prepare) | 0.31 | 0.51 | PO | DO | für | Mahlzeit (meal)<br>Empfang (reception)<br>Überraschung (surprise) |
| suchen<br>(search) | 0.36 | 0.45 | PO | PO | für | Aufgabe (task) |
|  |  |  |  |  |  | Geschenk (gift) |
|  |  |  |  |  |  | Haus (house) |
| schlachten<br>(slaughter) | 0.38 | 0.32 | PO | PO | für | Schwein (pig) |
|  |  |  |  |  |  | Ziege (goat) |
|  |  |  |  |  |  | Huhn (chicken) |
| bewachen<br>(guard) | 0.39 | 0.41 | PO | PO | für | Schatz (treasure) |
|  |  |  |  |  |  | Schloss (castle) |
|  |  |  |  |  |  | Gefangenen<br>(prisoner) |
| zahlen (pay) | 0.41 | 0.55 | PO | DO | für | Eintritt (entrance<br>fee) |
|  |  |  |  |  |  | Gebühr (fee) |
|  |  |  |  |  |  | Lösegeld<br>(ransom) |
| deuten<br>(interpret) | 0.42 | 0.47 | PO | PO | für | Wetter (weather) |
|  |  |  |  |  |  | Traum (dream) |
|  |  |  |  |  |  | Zeichen (sign) |
| reservieren<br>(book) | 0.43 | 0.45 | PO | PO | für | Tisch (table) |
|  |  |  |  |  |  | Ticket (ticket) |
|  |  |  |  |  |  | Platz (place) |
| verkaufen<br>(sell) | 0.45 | 0.7 | PO | DO | an | Goldfisch<br>(goldfish) |
|  |  |  |  |  |  | Blume (flower) |
|  |  |  |  |  |  | Buch (book) |
| liefern<br>(deliver) | 0.55 | 0.55 | DO | DO | an | Paket (package) |
|  |  |  |  |  |  | Vorrat (supplies) |
|  |  |  |  |  |  | Essen (food) |
| übergeben<br>(hand over) | 0.61 | 0.8 | DO | DO | an | Belohnung<br>(reward) |
|  |  |  |  |  |  | Rose (rose) |
|  |  |  |  |  |  | Rechnung (bill) |
| reparieren<br>(repair) | 0.63 | 0.42 | DO | PO | für | Dach (roof) |
|  |  |  |  |  |  | Waschmaschine<br>(washing<br>machine) |
|  |  |  |  |  |  | Auto (car) |
| leihen (lend) | 0.64 | 0.96 | DO | DO | an | Fahrrad (bike) |
|  |  |  |  |  |  | Jacke (coat) |
|  |  |  |  |  |  | Schirm<br>(umbrella) |
| verabreichen<br>(administer) | 0.68 | 0.98 | DO | DO | an | Tablette (pill) |
|  |  |  |  |  |  | Medizin<br>(medicine) |
|  |  |  |  |  |  | Spritze (injection) |
| reichen<br>(hand sth to<br>so.) | 0.69 | 0.92 | DO | DO | an | Koffer (suitcase) |
|  |  |  |  |  |  | Salz (salt) |
|  |  |  |  |  |  | Dose (can) |
| servieren | 0.72 | 0.71 | DO | DO | für | Kaffee (coffee) |
| (serve) |  |  |  |  |  | Wein (wine) |
|  |  |  |  |  |  | Speise (dish) |
| beschreiben<br>(describe) | 0.74 | 0.78 | DO | DO | für | Problem<br>(problem) |
|  |  |  |  |  |  | Spiel (game) |
|  |  |  |  |  |  | Aussicht (view) |

Previous research has suggested that in the brain, sentence-level language processing activates a widespread bilateral but left-dominant network of inferior frontal and middle and superior temporal regions (spanning from anterior to posterior areas) (Friederici & Gierhan, 2013; Hagoort, 2014; Hagoort & Indefrey, 2014). More specifically, these areas have been shown to be involved in processing PO and DO structure and distinguish between these as shown by pattern classification (Allen, Pereira, Botvinick, & Goldberg, 2012). Syntactic processing in particular seems to be guided by two key areas in left inferior frontal and left posterior middle temporal gyrus (Segaert, Menenti, Weber, Petersson, & Hagoort, 2012). The regions of the language network are highly interconnected. The arcuate fasciculus connects inferior frontal with the posterior middle/superior temporal gyrus (Catani, Jones, & Ffytche, 2005; Friederici, 2009) and the uncinate (in connection with the inferior fasicle) connects the temporal pole with the inferior frontal lobe via a more ventral route in the brain.

In recent years several studies have investigated the neural networks underlying predictive influences on language processing using a variety of different linguistic information (syntax: (Bonhage, Mueller, Friederici, & Fiebach, 2015; Henderson, Choi, Lowder, & Ferreira, 2016), words: (Willems, Frank, Nijhof, Hagoort, & Van den Bosch, 2015), semantics: (Lau, Weber, Gramfort, Hämäläinen, & Kuperberg, 2016; Weber, Lau, Stillerman, & Kuperberg, 2016) and speech: e.g. (Holdgraf et al., 2016)). These have uncovered predictive influences on processing within the areas related to processing the linguistic information (Bonhage et al., 2015; Henderson et al., 2016; Lau et al., 2016; Weber et al., 2016) as well as influences of areas that are not at the core of the language networks, such as the anterior cingulate (ACC) and subcortical structures (Bonhage et al., 2015; Weber et al., 2016). In particular, networks involved in cognitive control and adaptation (Botvinick, Cohen, & Carter, 2004; Shenhav, Cohen, & Botvinick, 2016) are likely to modulate areas related to processing the linguistic information, such as left inferior frontal gyrus (LIFG) and left middle/superior temporal gyrus (LM/STG), depending on e.g. the predictive validity of the input (Weber et al., 2016). The study by Henderson and colleagues (2016) found that the left inferior frontal gyrus and left anterior temporal lobe regions showed ‘syntactic surprisal’ effects, a measure of predictability of a given word’s syntactic category given its preceding context. In general, ‘surprisal’ is used as a measure in studies on prediction to quantify how unexpected some information is given the previous context. A high level of surprisal thus indicates the violation of a prediction. In another study on prediction in language processing, Weber and colleagues (2016) investigated how the statistics of the input, the proportion of semantically related to unrelated pairs of words between blocks, influences semantic processing and found enhanced LIFG to ACC connectivity under conditions of higher predictive validity. Modulations in the statistics of the input thus lead to a change in coupling between the language network and regions related to cognitive control, changing information flow when the input was more predictable. Furthermore, the predictive validity of the input (proportion differences between blocks) modulated the semantic priming effect within the language network, with a stronger priming effect (hemodynamic response suppression) in case of higher predictive validity. Given this prior work we assume that a large-scale network involving language and cognitive control regions is involved in using the linguistic context to modulate language processing. Accessing linguistic information such as the mental representation of words from memory will lead to a local expectation of which structure this word is likely to be paired with and will lead to a modulation of processing within the language network. On the other hand, the ACC is involved in keeping track of the frequencies in the input leading to expectations regarding words and structures more globally.

In the current experiment we were thus interested how different types of information that could be used for prediction modulate how the brain processes sentence structures. More specifically, we wanted to know whether these different types of information integrated on different time-scales (over the experiment versus around a word) would recruit different neural networks when used to modulate language processing. We expected the ACC and areas related to cognitive control to be responsive when the statistics of the linguistic environment are manipulated (as in (Weber et al., 2016)) and the core language network to be sensitive to surprisal based on live-long experience with a language (such as expectations of a certain syntactic category as in (Henderson et al., 2016)). That different types of information can have different effects on the neural processing of sentences is also underlined by different types of context leading to different types of ERP effects in studies manipulating local and global context on the semantic and discourse level (e.g. (Boudewyn, Long, & Swaab, 2015; Brothers, Swaab, & Traxler, 2015)). Here, we thus manipulated the statistics of the language input, namely the current distribution of sentence structures in a block, as well as using biases that were learned throughout the experience with a language, namely verb preferences. Participants read sentences with prepositional object and double object structures. The verbs used had a preference for one or the other structure in everyday language use (the syntactic preference of the verb could thus be used to predict which syntactic structure was likely to come up next). Moreover, we had three different blocks of sentences with different proportions of prepositional object (PO) and double-object (DO) sentences (Balanced Distribution: 50% DO/50% PO; Unexpected Distribution: 25% DO/75% PO; Expected Distribution: 75% DO/25% PO; the unexpected distribution is unexpected in the light of the DO structure being generally more frequent in German). This manipulation of the input statistics would make one structure more likely to come up next within a certain block. While participants were reading sentences, we acquired functional MR images, to investigate the underlying neural networks in prediction. Our hypotheses were:

1. Regions related to sentence-level/syntactic processing in the brain, specifically the LIFG and posterior LM/STG (the prediction that these two regions in particular will show these effects are based on neuroimaging studies of syntactic priming (Schoot, Menenti, Hagoort, & Segaert, 2014; Segaert et al., 2012, 2012; Weber & Indefrey, 2009) and a recent meta-analysis of sentence-level processing (Hagoort & Indefrey, 2014)) change their activation levels in response to verb specific syntactic surprisal, with larger surprisal leading to increased activation.
2. Changes to the current statistical environment, the relative frequency, of syntactic structures will lead to adaptations both within the sentence processing network as well as areas related to cognitive control, such as the ACC that monitors the statistical contingencies of the input. The unexpected distribution of statistical structures should engage the ACC the most with higher activations for the currently infrequent type of structure.
3. These regions outside the language network will interact with regions in the language network to adapt to the nature of the language input. These connectivity patterns should follow the pattern described under 2).
4. The two types of predictive information might interact. We expect the interaction to occur within the language network. If based on both the verb-based preference and the current environment a certain structure is expected but another shown, this surprisal effect might be stronger than if the surprisal is based on the verb-based preference alone.

## Materials and Methods

### Participants

We tested 21 German native speakers (7 male) and excluded one (male) participant from further analyses due to technical issues during acquisition. Behavioural responses were not recorded in the logfile of one subject due to a technical malfunction and were thus not included in the behavioural analysis. However, as online monitoring of the subject during the experiment had indicated task engagement this participant was kept in the fMRI analysis.

All participants were right-handed (as assessed by a German version of the Edinburgh Handedness Inventory (Oldfield, 1971), had normal or corrected-to-normal vision and no history of neurological impairments. The participants received compensation for their participation in the experiment and gave written informed consent before the study started. The study was approved by the internal review board of Carl von Ossietzky University Oldenburg in accordance with the declaration of Helsinki.

### Stimuli and Design

The experimental stimuli consisted of German ditransitive sentences, (i.e. sentences with verbs taking two arguments) half of them were double-object constructions (DO), half prepositional object ones (PO). The agents and patients in the sentences were always ‘Frau’ (woman), ‘Mann’ (man), ‘Kind’ (child). The theme (the other argument) varied to fit the verb (3 different potential themes per verb; see Table 1 for a list of verbs and nouns). Several ideas for themes were taken from Segaert and colleagues (2014) and Loebell and Bock (2003). The 8 ditransitive verbs were chosen so that they could occur both in the double-object and the prepositional construction (see Table 1 for a list of the different verbs, their themes and prepositions; see the introduction for example PO and DO sentences). Half of the verbs had a preference (a greater likelihood of being paired with a certain structure) for the double-object construction, half for the prepositional object construction.

The preference values were based on pretest rating results in 42 German native speakers of a previous study (Segaert et al., 2014). We made sure that the length of the two sets of verbs (PO-preference and DO-preference verbs) did not differ in length from each other (p=.47) and that their log lexical frequency values matched (based on subtlex, p=.33 (Marc Brysbaert et al., 2011) using independent two-sample t-tests. Additionally, we acquired data on the participants’ individual verb preferences in a post-test one week after the main experiment. This post-test consisted of sentence completions (“The woman builds…”), 3 utterances for each verb, and counted the number of PO and DO completions per verb to get a participants’ verb preference values. The preference values from the previous study were used to for the initial categorisation into PO and DO preference verbs. However, we used the group preference values from the post-test in current study for the analysis as we assume that these values more accurately reflect the biases of the investigated group of participants.

The structure of the stimuli was as follows. The experiment consisted of three blocks of sentences with different statistics. In the first block DO and PO constructions occurred equally often. This block was included as a potential baseline to investigate DO and PO sentence processing when they were equally probable. In the second block we changed the proportion of the two types of sentence structures. DO sentences occurred 75% of the time and PO sentences 25% of the time, this block is similar to everyday language use in German. The third and last block switched the proportions of DO and PO sentences, now 25% had a DO sentence structure and 75% a PO structure, this block is less similar to everyday language use in German. The order of the last two blocks was counterbalanced across participants. Each sentence structure occurred with equal amounts of DO and PO-preference verbs. Within each block there were 4 conditions: DO structure with DO verb-preference; DO structure with PO verb-preference; PO structure with DO verb-preference and PO structure with PO verb-preference. Each condition contained at least 24 sentences. Thus, in the block with equal probability for each structure, each condition (bias towards PO – PO structure; bias towards PO – DO structure; bias towards DO – PO structure; bias towards DO – DO structure) contained 24 items. In blocks where one of the structures occurred 75% of the time, the conditions containing the more frequent structure included 72 sentences and the infrequent conditions 24 sentences.

Participants were instructed to read the sentences carefully and silently in their head. Randomly interspersed, after on average 8 sentences (after 12% of the sentences) a comprehension question (e.g. “Was the previous sentence about a child?” or “Did the man buy the boat?”) was asked and the participant was instructed to press one of two buttons for yes or no.

In summary we constructed a design with three factors ‘Structure Statistics (50% DO; 75% DO; 25% DO), ‘Verb Preference’ (bias towards PO; bias towards DO structure) and ‘Structure’ (PO or DO).

We also designed a language network localiser task to obtain a group specific localisation of the language network. The task consisted of four conditions: sentences, random word lists, sentence-like lists of pseudo words and random pseudo word lists. The sentence condition consisted of 24 ditransitive sentences (12 DO, 12 PO) made up of different verbs and nouns compared to the main experiment. The random word lists condition was created by generating another set 24 dative sentences that were then scrambled within and across the sentences (which of the sentence lists was used for the random word lists was counterbalanced across participants). The sentence-like lists of pseudo words and random pseudo word lists were created by replacing the words in the previous two conditions with pseudo words that matched the real words in length and transitional probabilities using Wuggy (Keuleers & Brysbaert, 2010). During the sentence localiser task the different conditions were presented in random order. As in the main experiment the noun phrases (determiner and noun) of the sentences were presented together on the screen (and the other conditions followed this basic format). As for the main experiment, the participants were instructed to read the sentences and word lists attentively and silently.

## Experimental Procedure

In the MR scanner stimuli were visually presented to the participants via a mirror system. The sentences were presented in light grey font (font size 20; type Verdana) on a black background. Experimental trials were delivered in segments (i.e. noun phrases (e.g. ‘Der Mann’) were presented together). Each segment was displayed for 500ms followed by a 100ms blank screen. Between experimental trials a fixation cross was displayed on the screen. At random intervals, comprehension questions were asked after a sentence. This question was displayed for 4 seconds and participants pressed one of two buttons to answer the question with yes or no. This was again followed by a fixation cross. The duration of the fixation crosses, and thus the inter-trial interval, varied between .4 and 10s and was predetermined by a dedicated software (Dale, 1999) used to optimise the timing of trials to remove the overlap between trials from the hemodynamic response estimates.

### Structural and functional MRI data acquisition

Structural and functional magnetic resonance images were acquired using a 3T Siemens Verio scanner equipped with a 8-channel head coil. The functional volumes were acquired using an EPI sequence (30 axial slices (AC-PC aligned), 3.1×3.1 mm voxel size, repetition time=2s, echo time=30 ms, ascending acquisition). One dataset of T1-weighted high-resolution structural images (1mm isotropic voxel size, MPRAGE sequence) was acquired at the end of each session.

### Data analysis

Pre-processing as well as the first and second level analyses of the fMRI data made use of the SPM12 software (www.fil.ion.ucl.ac.uk/spm), a MATLAB based toolbox (www.mathworks.com/matlab). In particular:

### Preprocessing

The images were spatially realigned to the first image of the first block and then across blocks and then slice-time corrected. The functional images were co-registered to the structural image by co-registering the mean functional image to the structural MPRAGE. The anatomical image was segmented into grey and white matter and the spatial normalization parameters from the segmentation step were then used to normalize the functional images. Finally, the images were smoothed with an 8mm full width at half maximum (FWHM) Gaussian kernel.

### First Level: Localizer

We acquired a language localizer at the end of the fMRI epxeriment. Its design matrix consisted of one block with one regressor per experimental condition (sentences, random word lists, sentence-like lists of pseudo words and random pseudo word lists). The actual onset of the first segment of a sentence/word list was taken as the onset time of a trial and the actual duration of the event was modelled. In addition we added 6 movement regressors. Per subject we identified contrast images that were then taken to the second level for a random effects group analysis.

### First Level: Main Experiment - Activation

Per subject, the design matrix for the main part of the experiment consisted of three blocks, one per ‘Current Structure Statistics’ condition (Balanced: 50% DO; Unexpected Distribution: 25% DO; Expected Distribution: 75% DO). Within each block we modelled event-related regressors for each of the conditions of the factor ‘Structure’: i.e. DO and PO sentences. The number of trials used in those regressors was kept constant. This meant that for the 25% DO block, the remaining (randomly selected) PO sentences went into a separate ‘filler’ regressor. Also in the 75% DO block the additional DO sentences went into a separate ‘filler’ regressor. For each of the sentence regressors of the factor ‘Structure’ we added another regressor, a parametric modulator, reflecting verb-based syntactic surprisal (‘Verb-based syntactic surprisal’: which we defined as the negative log probability of encountering a syntactic structure given the verb-preference, a larger value indicates an unexpected, surprising event) of each PO and DO sentence. Thus, if a verb had a strong bias towards a PO structure but a DO structure was shown, the surprisal value would be high. The verb-preference values were based on the post-test results of the current group of participants. These reflect the biases of the current group of participants (compared to the questionnaire values from a separate group of participants that we based our initial verb selection on; as Table 1 shows, the values for the original questionnaire, column 2, and the post-test from the present group, column 3, are largely in the same direction with a couple of deviations). We investigate both the main effects of ‘verb-based syntactic surprisal’ as well as its effect per sentence structure (PO and DO) as their overall different distribution might influence verb-based syntactic surprisal effects. As for the localizer, the onset of the first segment was taken as the time of onset, and the actual duration of the sentence was modelled. In addition we added 6 movement regressors. Per subject we identified contrast images that were then taken to the second level for a random effects group analysis. For the analysis of the interaction between ‘Structure’ and ‘Current Structure Statistics’ these were the contrast images of the regressors per structure (PO or DO) per ‘Current Structure Statistics’ block against the implicit baseline. For the analysis of ‘verb-based syntactic surprisal’ these were the contrast images of parametric modulation regressors (per structure and block) against the implicit baseline.

### First Level: Main Experiment - Connectivity

Task-related functional connectivity analyses were carried out using the generalized context-dependent psychophysiological interactions (gPPI) toolbox (McLaren, Ries, Xu, & Johnson, 2012). As a seed region we chose the expected ACC activation from the interaction between ‘Current Structure Statistics’ and ‘Structure’ (voxel-threshold p<.001, cluster-level p_FWE_<.05). The time series of the seed region was added as an explanatory variable to the model. We modelled regressors describing the connectivity from the seed for all conditions described for the main activation analysis above (main regressors and parametric modulators), as well as regressors corresponding to the activity in each of the experimental conditions.

### Second-level Analysis - Localizer

We built a flexible factorial design with a regressor per experimental condition (sentences, random word lists, sentence-like lists of pseudo words and random pseudo word lists) as well as regressors to model the within subject-effect (thus one regressor per subject).

### Second-level Analysis - Main Experiment: Activation Analysis

We built two different design matrices to look at the group-level results of the main experiment. The first one was based on the two sentence regressors (PO structures; DO structures) per block and was designed to investigate the modulation of the processing of sentences by the surrounding syntactic statistics. This analysis focused on the ‘Unexpected Distribution: 25% DO structure’ and the ‘Expected Distribution: 75% DO structure’ blocks because the position of the ‘Balanced Distribution: 50% DO structure block’ (which was designed as a baseline measurement) was not counterbalanced across subjects. The flexible factorial designs were built using the factors ‘Subject’ (20 regressors, one regressor per subject, to model within-subject effects), ‘Structure’ (PO structures; DO structures) and ‘Current Structure Statistics’ (the different blocks), with one regressor per condition.

The second design matrix had the same design setup but was based on the parametric modulators based on verb-based syntactic surprisal values for the two types of structures per block. This design matrix was designed to look at the effect of verb-based syntactic surprisal overall (across all 3 blocks) per type of structure (factor “Structure) and its interaction with the syntactic statistics (factor ‘Current Structure Statistics’). As in the other model we also included the factor ‘Subject’ to model within-subject effects.

### Second-level Analysis - Main Experiment: Functional Connectivity Analysis

For the task-related connectivity analysis, we evaluated a design matrix similar to the one for the activation analysis, but based on the PPI regressors (McLaren et al., 2012). This analysis focused on the interaction between ‘Current Structure Statistics’ and ‘Structure’ as we wanted to look at the interaction between language and non-language regions for this contrast. The seed region was defined based on the interaction between ‘Current Structure Statistics’ and ‘Structure’ in the activation analysis to see with which regions the region showing a modulation by the current linguistic environment interacted.

For all analyses, we report effects at a voxel-level threshold of p<.001 and a cluster extent threshold of 25 voxels to show patterns and trends. For statistical inference we highlight those activations that reach a cluster-level FWE-corrected threshold of p<.05 or Small Volume Correction (Worsley et al., 1996) at the peak at p<.05. As we expected effects to be located in the canonical language network we used Small Volume Correction (SVC) with the left-hemisphere regions defined in the localizer (see highlighted activations in Table 2) where appropriate. All reported coordinates are in MNI space.

**Table 2.**
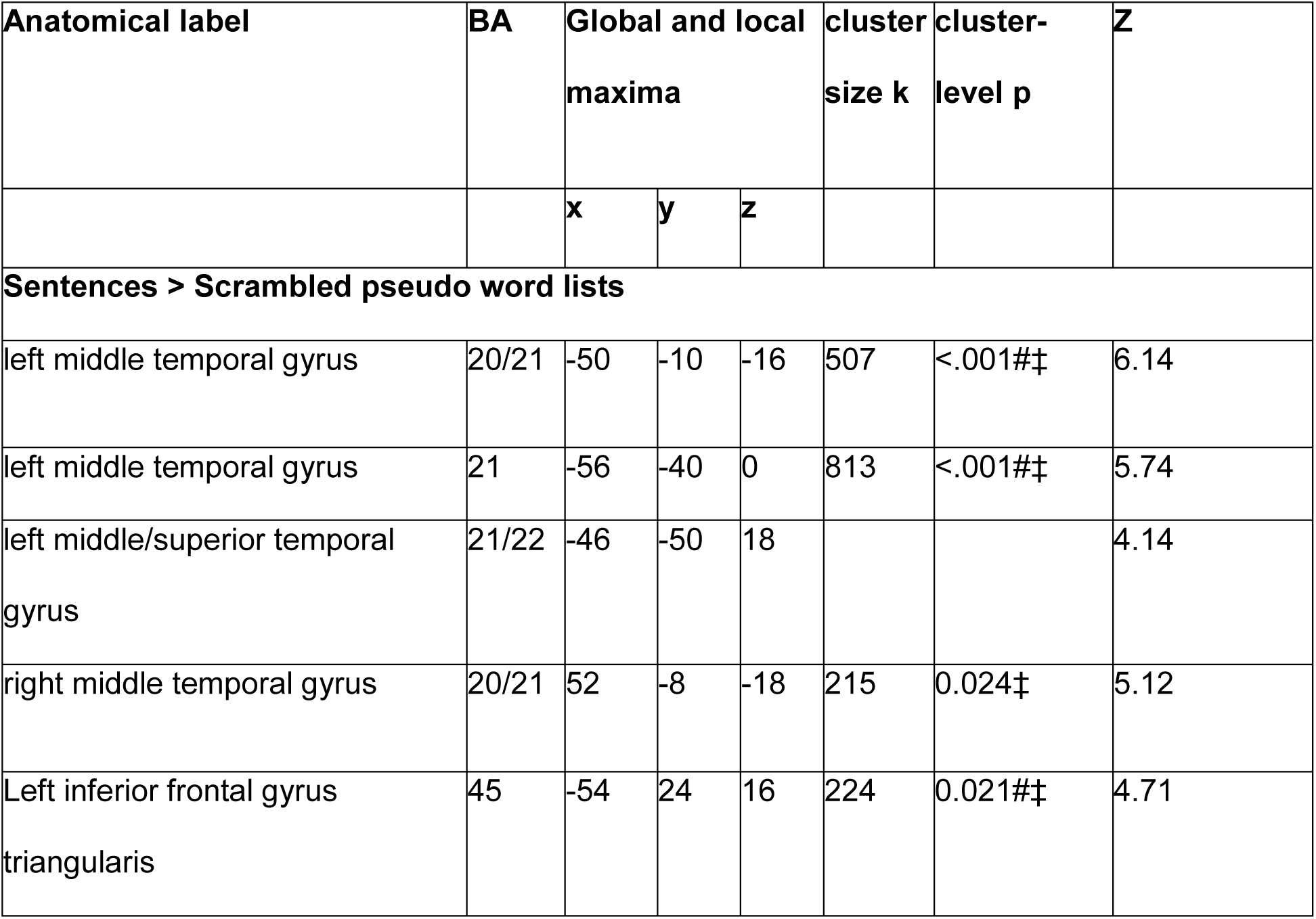
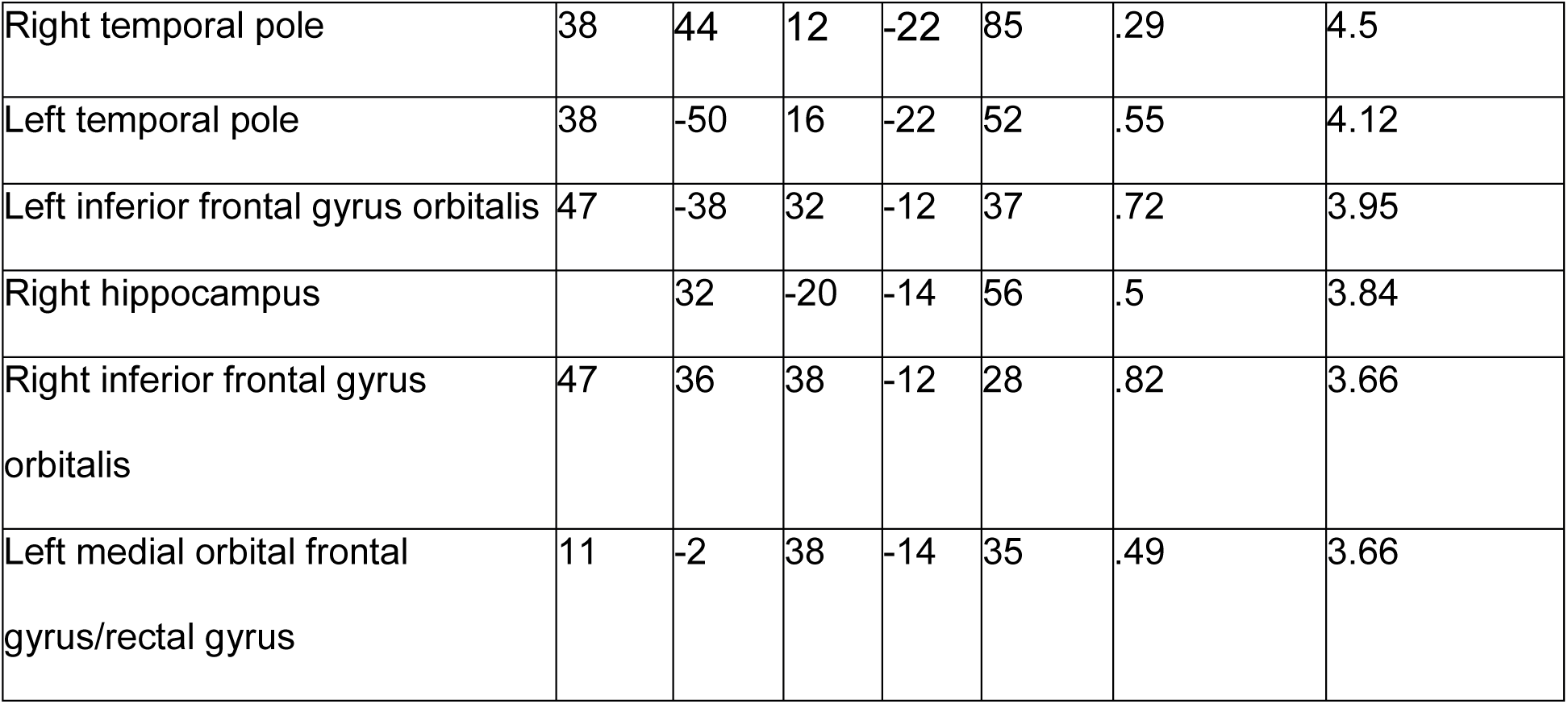
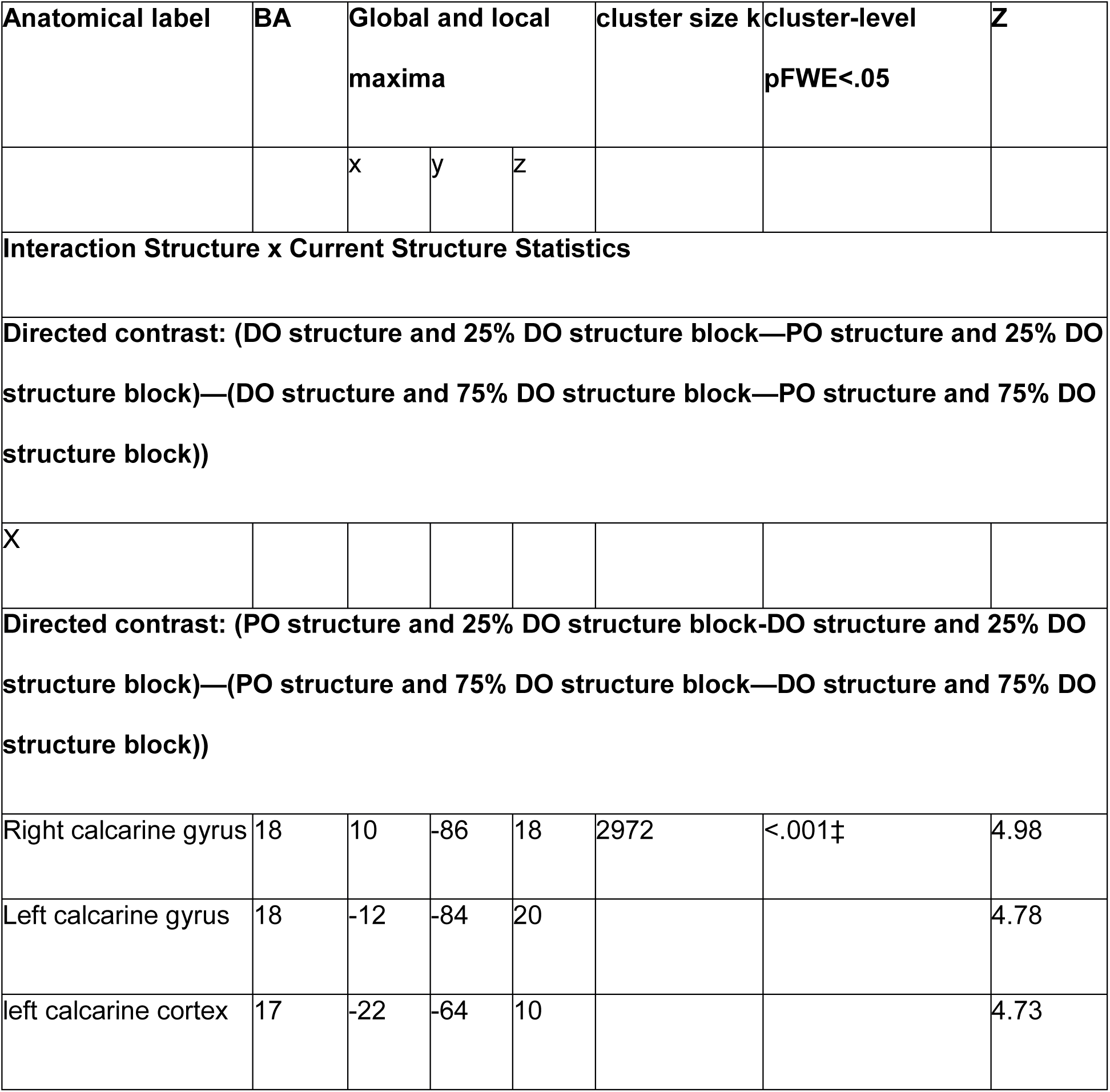
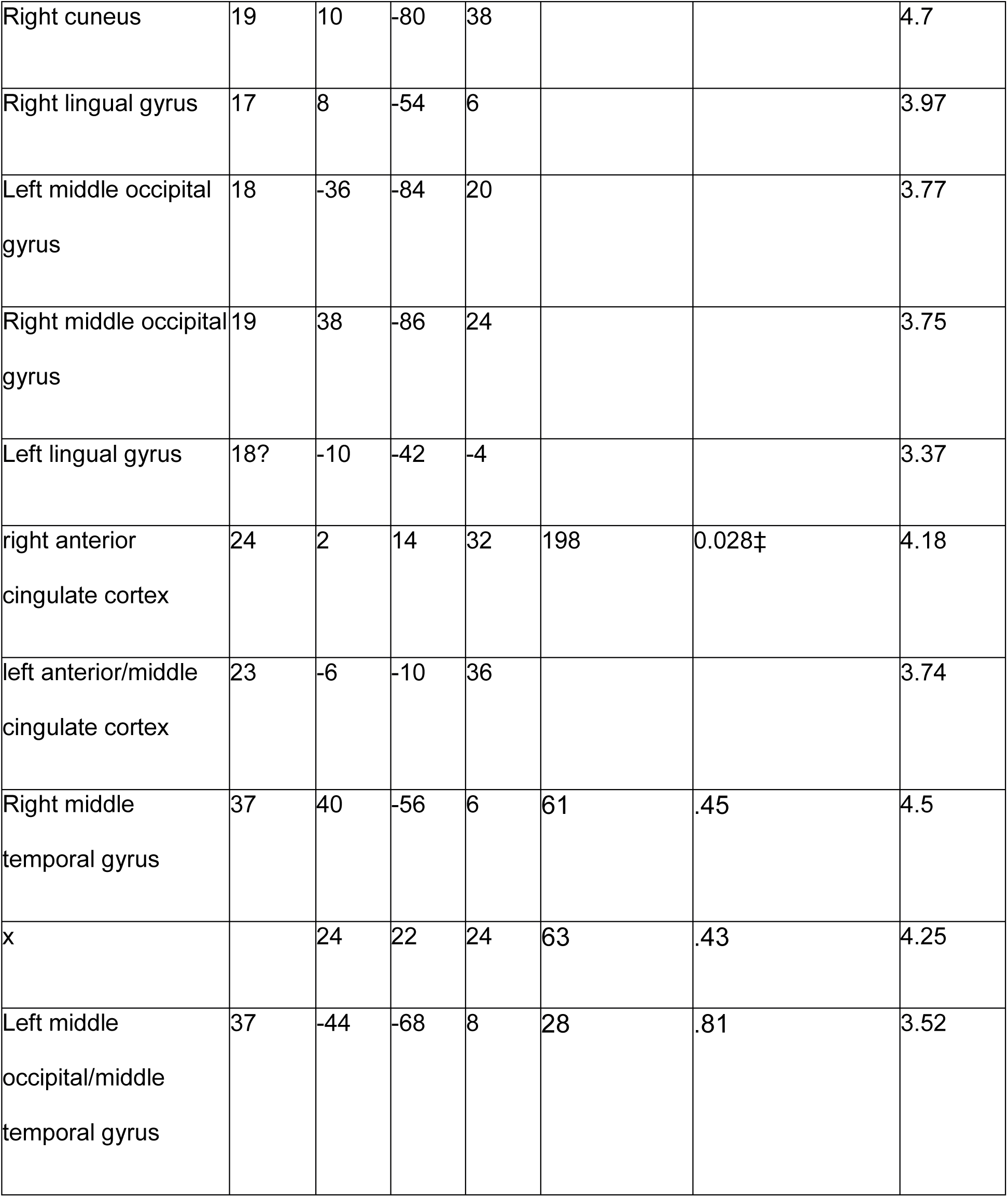
Whole-brain activations for the activation effects for sentence structures. Listed are local maxima more than 20mm apart. All clusters at a voxel-level threshold of p<.001, k=25 are reported, those that reach cluster-level FWE correction or small volume correction are marked by ‡.

## Results

We will firstly briefly describe the behavioural results, i.e. the performance on the questions during the experiment and the post-experimental questionnaire. The results of the localizer will serve both as a sanity check showing that a canonical language network is activated in our participants and to define regions of interests that will be used as seed regions in the connectivity analysis as well as for small volume correction.

Next, we will describe the effects of the current context (‘Current Structure Statistics’) on the processing of PO and DO sentence structures (‘Structure’). This will thus characterize the interaction between ‘Current Structure Statistics’ and ‘Structure’, both for the activation and the connectivity analysis. This will be followed by an investigation of the effect of the parametric modulator ‘Verb-based syntactic surprisal’ (and planned comparisons of the effect of ‘Verb-based syntactic surprisal’ per sentence structure, DO or PO) and their interactions with the current context (‘Current Structure Statistics’).

### Performance Questions during the Experiment

On average participants got 91% of the questions correct (range=80-100%, SD=7%), showing that they paid attention to the meaning of the sentences while reading.

### Performance Post-test

Column three of Table 1 illustrates the verb-preference values based on the post-test. Values from 3 participants were not included in these group averages because they did not return the questionnaire (2 participants) or did not fill in the questionnaire with any ditransitive sentences as answers (1 participant). Two participants did not fill in any ditransitive sentences for 4 and 5 of the verbs respectively and these missing cells were replaced with the group average values for these verbs.

### Localizer

The contrast of sentences versus scrambled pseudo word lists (a complex visual baseline) revealed activation in a canonical language network including, left inferior frontal gyrus, left middle and superior temporal gyrus as well as the right middle temporal gyrus (see Table 2). The activation results in the left hemisphere were used to define regions of interests for the main experiment as well as for small volume correction (Worsley et al., 1996). We chose this contrast as it should capture regions involved in syntactic, semantic and lexical processing.

Table 2. Whole-brain activations for the language localizer task. Listed are local maxima more than 20mm apart. All clusters at a voxel-level threshold of p<.001, k=25 are reported, those that reach cluster-level FWE correction or small volume correction are marked by ‡. Clusters used for Small Volume Correction for the main experiment are marked by #.

### Activation - Interactions between ‘Current Structure Statistics’ and ‘Sentence Structure’

We found effects in a cluster spanning cuneus, precuneus and occipital regions as well as a cluster in left and right anterior/middle cingulate cortex, for the interaction between ‘Structure’ and ‘Current Structure Statistics’, see Table 2 and Figure 1. Follow-up t-test revealed that the activation pattern within the cingulate cortex cluster differed between the PO and DO structure in the ‘Unexpected Distribution’ block (PO more frequent than DO sentences) with a larger activation for the PO structure, t(19)=-3.2, p=.005 as well as in the ‘Expected Distribution’ (DO more frequent than PO sentences) block (t(19)=2.6 p=.02), albeit in opposite direction, the DO structure showing higher activation than the PO structure. One of the contributing factors might be a change in activation patterns for the PO structure across blocks, there was a trend towards a higher activation for the PO structure in the ‘Unexpected Distribution’ t(19)=1.8, p=0.09, while the activation pattern for the DO structure staid the same across blocks t(19)<1.

**Figure 1.**
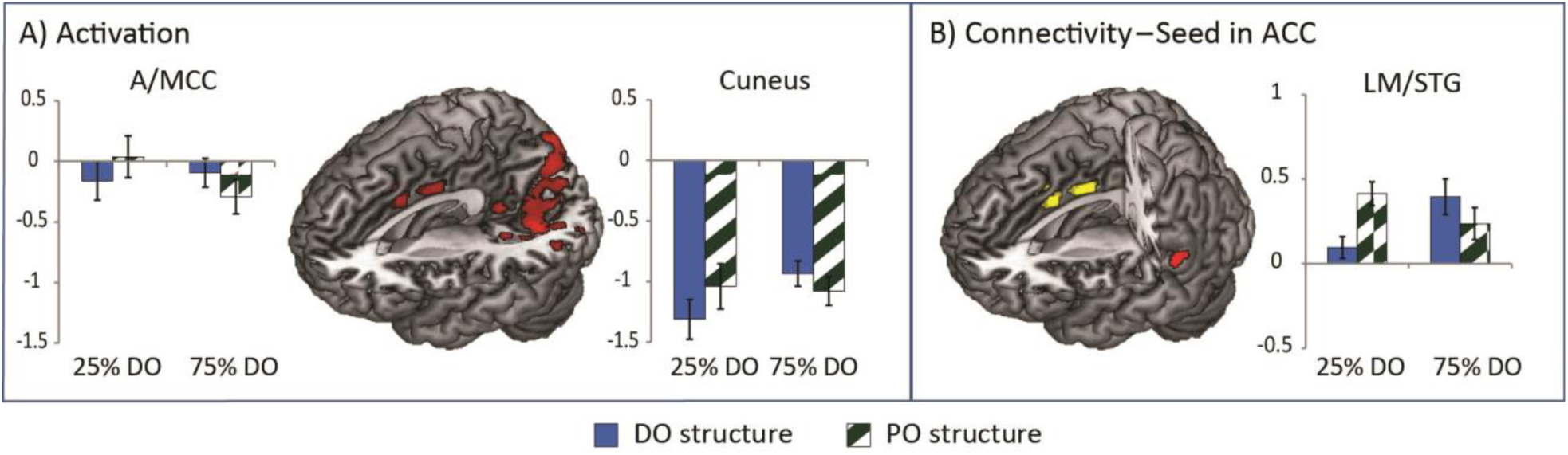
Interactions between type of structure (DO vs. PO) and ‘Current Structure Statistics’ (25% DO structure: “Unexpected Distribution” vs 75% DO structure: “Expected Distribution”). A) whole-brain activation results, B) PPI connectivity results from a seed in ACC (in yellow). Effects are shown at a voxel-level significance threshold of p<.001 with a cluster-level threshold pFWE<.05 or pSVC<.05. Bar graphs show mean contrast values per condition for a cluster. See Table 2 for a complete list of activations and connectivity patterns.

The follow-up t-tests for the cuneus/occipital cluster showed a lower activation for the DO than for the PO structure in the ‘Unexpected Distribution’ block: t(19)=-4.5, p<.001, and the opposite pattern in the ‘Expected Distribution’ block: t(19)=2.5, p=.02. Here, the DO structure changed activation patterns across blocks, t(19)=-2.1, p=.048, while the PO structure did not t(19)<1.

### Connectivity - Interactions between ‘Current Structure Statistics’ and ‘Sentence Structure’

We found task-related functional connectivity patterns from the seed in ACC to the left middle/superior temporal gyrus, see Table 3 and Figure 1 B. This connectivity pattern was driven by a difference in the strength of this connection between the DO and PO structure in the ‘Unexpected Distribution’ block, t(1,19)=-4.6, p<.001, whereas there was only a weak trend towards a difference in the ‘Expected Distribution’ block, t(1,19)=1.8, p=.09. This was driven both by an increase in connectivity for the DO structure from the unexpected to the expected distribution, t(19)=-3.1, p=.006 as well as a trend towards the opposite pattern for the PO structure, t(19)=1.9, p=.066.

**Table 3.**
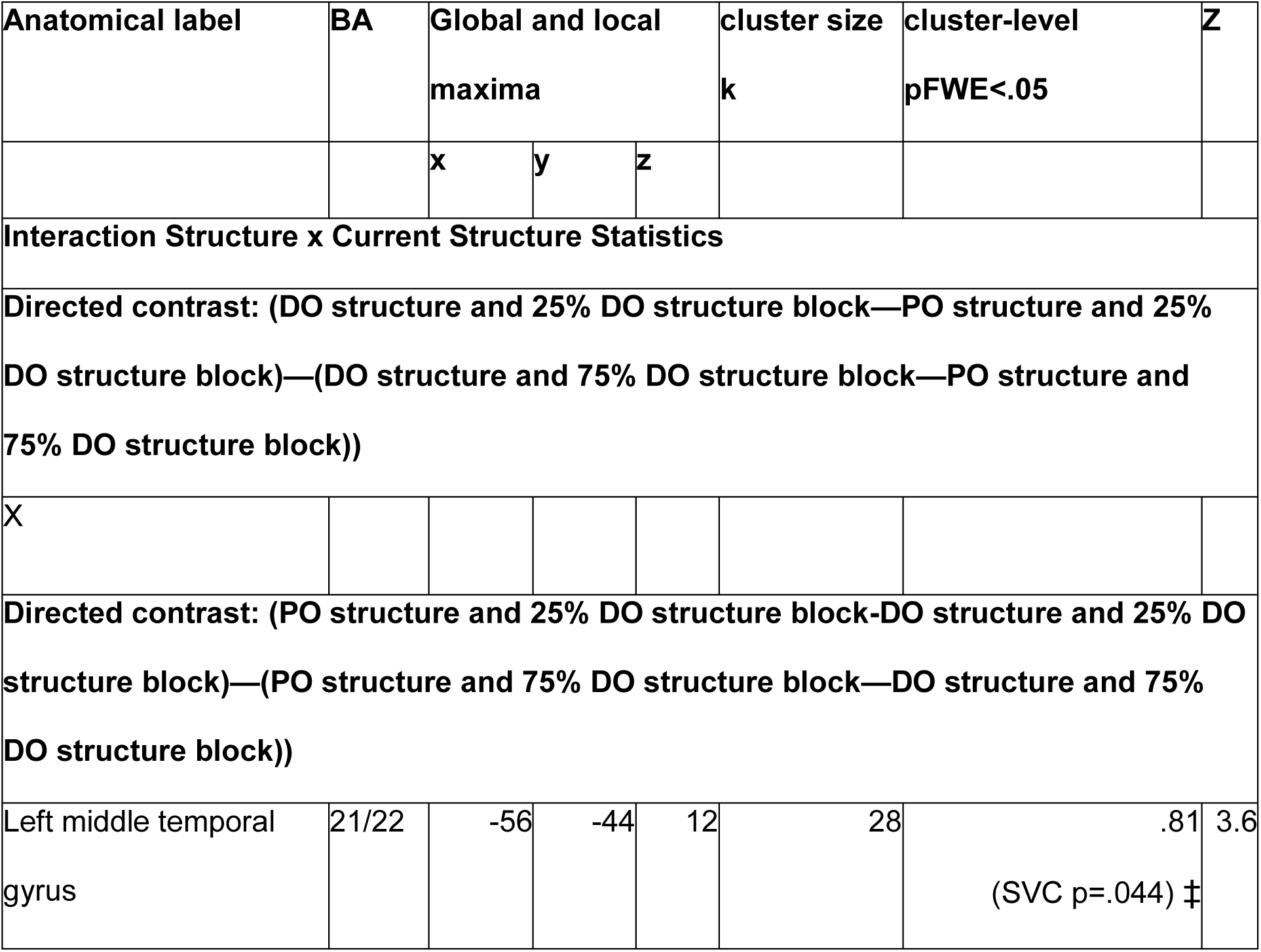
Whole-brain effects for the connectivity results for sentence structures. Listed are local maxima more than 20mm apart. All clusters at a voxel-level threshold of p<.001, k=25 are reported, those that reach cluster-level FWE correction or small volume correction are marked by ‡.

| Anatomical label | BA | Global and local maxima |  |  | cluster size k | cluster-level pFWE<.05 | Z |
| --- | --- | --- | --- | --- | --- | --- | --- |
|  |  | x | y | z |  |  |  |
| **Interaction Structure x Current Structure Statistics** |  |  |  |  |  |  |  |
| **Directed contrast: (DO structure and 25% DO structure block—PO structure and 25% DO structure block)—(DO structure and 75% DO structure block—PO structure and 75% DO structure block))** |  |  |  |  |  |  |  |
| X |  |  |  |  |  |  |  |
| **Directed contrast: (PO structure and 25% DO structure block-DO structure and 25% DO structure block)—(PO structure and 75% DO structure block—DO structure and 75% DO structure block))** |  |  |  |  |  |  |  |
| Left middle temporal gyrus | 21/22 | -56 | -44 | 12 | 28 | .81<br>(SVC p=.044) ‡ | 3.6 |

### Activation - Main Effect ‘Verb-based Syntactic Surprisal’ and Interaction with ‘Current Structure Statistics’

Over all 3 blocks, no main effect of verb-based syntactic surprisal (the negative log probability of encountering a syntactic structure given the verb-preference) was found, two clusters in LM/STG and Precuneus did not survive cluster-level correction. There was also no interaction of syntactic surprisal with structure or ‘Current Structure Statistics’. However, planned comparison of the verb-based syntactic surprisal effects, separately for PO and DO structures revealed a syntactic surprisal effect for the DO structure only. Regions in LIFG and LS/MTG (small volume corrected with regions of interests, see Table 2) and precuneus show higher activations with higher surprisal values (i.e. higher activation if the verb biased towards a PO structure, the structure that is generally encountered less frequently in German, but a DO structure was presented). See Table 4 and Figure 2 for effects of the parametric modulation of verb-based syntactic surprisal.

**Table 4.**
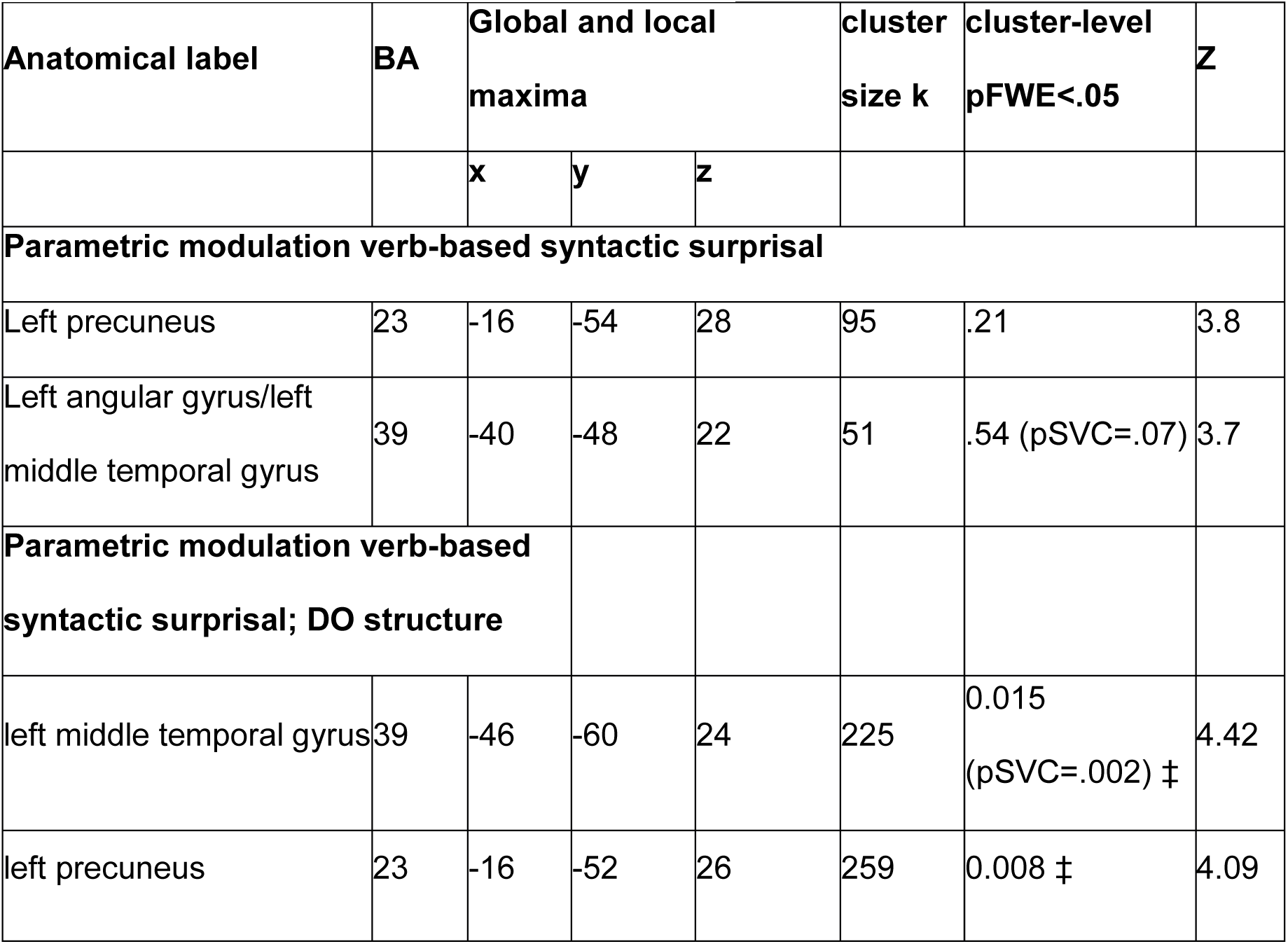
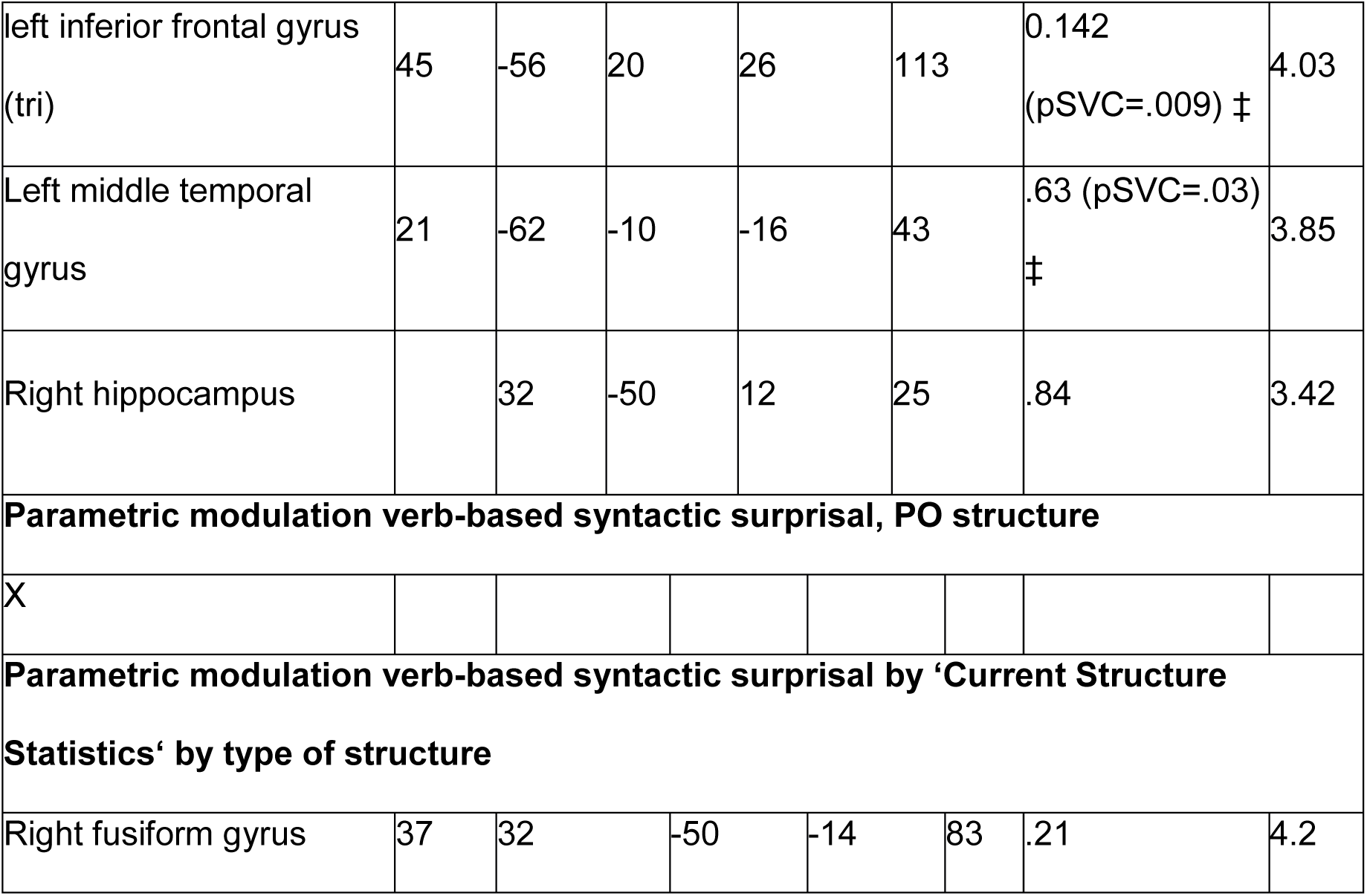
Whole-brain effects of the parametric modulations of verb-based syntactic surprisal. Listed are local maxima more than 20mm apart. All clusters at a voxel-level threshold of p<.001, k=25 are reported, those that reach cluster-level FWE correction or small volume correction p<.05 are marked by ‡.

| Anatomical label | BA | Global and local maxima |  |  | cluster size k | cluster-level pFWE<.05 | Z |
| --- | --- | --- | --- | --- | --- | --- | --- |
|  |  | x | y | z |  |  |  |
| Parametric modulation verb-based syntactic surprisal |  |  |  |  |  |  |  |
| Left precuneus | 23 | -16 | -54 | 28 | 95 | .21 | 3.8 |
| Left angular gyrus/left middle temporal gyrus | 39 | -40 | -48 | 22 | 51 | .54 (pSVC=.07) | 3.7 |
| Parametric modulation verb-based syntactic surprisal; DO structure |  |  |  |  |  |  |  |
| left middle temporal gyrus | 39 | -46 | -60 | 24 | 225 | 0.015 (pSVC=.002) ‡ | 4.42 |
| left precuneus | 23 | -16 | -52 | 26 | 259 | 0.008 ‡ | 4.09 |
| left inferior frontal gyrus<br>(tri) | 45 | -56 | 20 | 26 | 113 | 0.142<br>(pSVC=.009) ‡ | 4.03 |
| Left middle temporal<br>gyrus | 21 | -62 | -10 | -16 | 43 | .63 (pSVC=.03)<br>‡ | 3.85 |
| Right hippocampus |  | 32 | -50 | 12 | 25 | .84 | 3.42 |
| **Parametric modulation verb-based syntactic surprisal, PO structure** |  |  |  |  |  |  |  |
| X |  |  |  |  |  |  |  |
| **Parametric modulation verb-based syntactic surprisal by 'Current Structure**<br>**Statistics' by type of structure** |  |  |  |  |  |  |  |
| Right fusiform gyrus | 37 | 32 | -50 | -14 | 83 | .21 | 4.2 |

**Figure 2.**
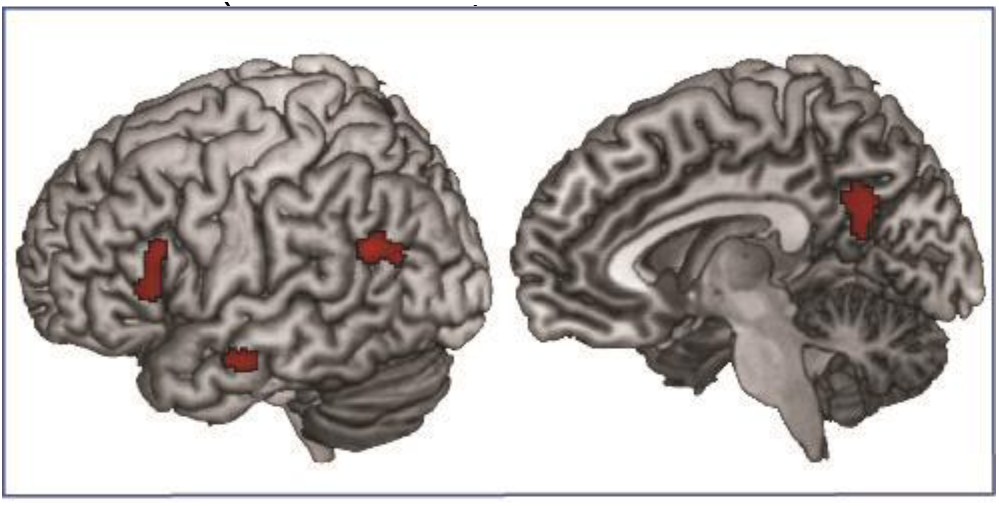
Parametric modulations of verb-based syntactic surprisal for the DO sentence structure. Effects are shown at a voxel-level threshold of p<.001, k=25, and survive FWE or SVC correction (see Table 4).

## Discussion

In this study we manipulated two types of information that can be used for prediction in sentence-level processing: the within-experiment context, i.e. the statistics of syntactic structures in different blocks, and verb-based syntactic surprisal, i.e. the preference for a syntactic structure given a verb. The results showed that changing the syntactic statistics of the current linguistic environment (the proportion of PO versus DO structures) in a block, resulted in the largest difference between the PO and DO structures in the block with the unexpected statistical distribution that was opposite to the one encountered in everyday life. Here, the PO structure, frequently presented in the experimental block but generally infrequent in everyday life, showed the highest activation in the anterior/middle cingulate and an increased functional connectivity from this node to the posterior parts of the language network (LM/STG). Conversely, the second manipulation, surprisal to see a structure given a certain verb when a verb had a PO verb-bias but a DO structure was encountered, resulted in an increased activation within the language network (left inferior frontal and left middle/superior temporal gyrus) as well as in the precuneus. Interestingly, we did not find any interactions between the syntactic statistics of the current linguistic environment and verb based syntactic surprisal effects.

### The effects of current structure statistics on processing sentence structures

The anterior cingulate cortex is sensitive to statistical contingencies in the language input (Weber et al., 2016) and is part of a larger network involved in cognitive control and adaptation to changing, volatile environments. Furthermore, several studies have suggested a prominent role of this region in the processing of unpredicted and infrequent events in the input (Behrens, Woolrich, Walton, & Rushworth, 2007; Botvinick et al., 2004; Shenhav et al., 2016; Vassena, Holroyd, & Alexander, 2017).

Interestingly, here we find that the activation pattern is not exclusively driven by the currently infrequent event, as we had predicted, but by the event that is unexpectedly frequent in the experiment. More specifically, the PO structure that is generally infrequent in everyday life but is suddenly frequent in one of our statistical environment blocks generates increased ACC activation compared to the other conditions (see Figure 1). Thus, in the current experiment we show that it is thus not only the case that the cingulate marks events as unexpected based on the current input but in a combination of current input statistics and a lifetime of experience with the statistics of sentence structures.

Also the cuneus and adjacent occipital areas appear to be sensitive to these statistical contingencies. These areas are part of the default mode network (Utevsky, Smith, & Huettel, 2014) and less deactivation for the more frequent structure might be related to its prominence disengaging parts of the default mode network.

When looking at connectivity from the ACC, to investigate how this region monitors the statistical contingencies of the input and it is functionally connected with other cortical areas, we find a responsive region in the left S/MTG, one of the core hubs of the language networks (Hagoort, 2014). Again, this interaction on the connectivity values was driven by a larger difference between the DO and the PO structure in the unexpected distribution block. The ACC and the posterior temporal region were most tightly interconnected for the PO structure in this block, the currently frequent structure that is generally the infrequent one. This tighter functional coupling between the ACC and the language network might reflect a role of the ACC in using its analysis of the statistical contingencies based on the current input and prior knowledge to weigh information flow in the language network.

In sum, the ACC might be engaged in tracking the statistics of the input and in communicating this information to relevant language regions such as LM/STG. However, this is done in a more sophisticated manner than previously thought as it appears to combine information of the current statistical environment with information on statistics in the environment that we learned over a lifetime. As the exact nature of this interaction, with the ACC appearing to track the ‘unexpectedly frequent’ event, was unexpected, the effect of the current statistical environment on language processing should be investigated in more depth in future studies.

### The effects of verb-based syntactic surprisal

The main effect of verb based syntactic surprisal in the left posterior temporal gyrus and precuneus had weak statistical power and did not survive cluster-level multiple comparison correction. However, planned comparisons separating verb-based syntactic surprisal effects for the DO and the PO structure (as they have generally different frequencies to begin with) revealed verb-based syntactic surprisal effects for the DO structure in the language network (LIFG and left posterior temporal gyrus) as well as the precuneus (see Figure 2). Such an effect was restricted to the presentation of DO sentences and this could be due to the fact that this is the more common syntactic structure in German. Thus, if based on the verb one would predict the more infrequent PO structure, this might lead to a strong reversal of the general prediction of a DO structure. If then a DO structure is shown after all this might lead to a larger surprisal effect than if the verb had biased towards a DO (with no large changes in prediction levels) but a PO was encountered.

Larger activation to verb-based syntactic surprisal reflects higher activations for disconfirmed predictions regarding which sentence structure will occur. This might reflect predictions down to the level of the predicted types of words and engage areas related to syntactic processing. The left inferior frontal and posterior temporal regions are specifically involved in syntactic processing of sentences (Menenti, Gierhan, Segaert, & Hagoort, 2011; Rodd, Vitello, Woollams, & Adank, 2015; Schoot et al., 2014; Segaert, Kempen, Petersson, & Hagoort, 2013; Segaert et al., 2012) with a specific focus for the processing and retrieval of lexical-syntactic information in left middle temporal gyrus (Snijders et al., 2009). While not a typical language network region, the precuneus showed sensitivity to syntactic structure repetition in some of these studies (Schoot et al., 2014; Segaert et al., 2013) and also in a meta-analysis (Rodd et al., 2015) and could thus be seen as part of the syntactic processing network.

Another study looking at syntactic surprisal effects in fMRI (Henderson et al., 2016) also found effects in left inferior frontal and temporal regions (albeit more anterior) among other regions (putamen, insula, fusiform gyrus and diencephalon). However, in their study syntactic surprisal was calculated as the surprisal to see a word of certain syntactic category given the previous words, this is different from the current surprisal of seeing a certain syntactic structure given a verb. The verb based syntactic surprisal effect that we find in left posterior temporal gyrus might be driven by activation of syntactic information linked to the verb in this area (Snijders et al., 2009).

In short, regions related to sentence-level and syntactic processing show a verb-based syntactic surprisal effect if a strong initial prediction towards the generally more infrequent structure is disconfirmed.

### The absence of an interaction effect between the current syntactic statistics and verb-based syntactic surprisal

In the present study we do not find any evidence for an interaction between verb-based syntactic surprisal and current syntactic statistics. Thus, the more local effect of predicting which structure will appear given a certain verb and the more global effects of using the statistical information of the wider environment to predict upcoming sentence structures, seem to be independent and subserved by different mechanisms. Verb-based syntactic surprisal is contained within the language network where predictive effects arise based on information stored in the mental lexicon. On the other hand, areas related to cognitive control, in this case the ACC, are in communication with the language network to modulate processing based on the statistical contingencies.

However, with fMRI we can only look at the overall activation level for the entire sentence obfuscating certain time-specific effects. In the future, using electroencephalography to look at ERP effects, such as the N400, which is sensitive to predictive validity (Lau, Holcomb, & Kuperberg, 2013) during reading of the post-verbal noun, might shed further light on potential interactions between these effects.

Moreover, one further limitation of the study lies in its limited set of verbs (16 in total). Thus, while we clearly had a modulation of the activation based on verb-based syntactic surprisal, the limited number of verbs limits the generalization over items.

In sum, we show that verb-based syntactic surprisal is processed ‘locally’ within the language network while the within-experiment context, the statistics of the input changes ACC activation and connectivity. The ACC appears to mark sentence structures as unexpected based not on the current input alone, but in a combination of current input statistics and knowledge of the frequency of different structures learned over a lifetime. The functional coupling between the ACC and the language network might suggest that the ACC has a top-down regulatory role on the processing within the language network.

## Acknowledgments

We would like to thank the members of the Applied Neurocognitive Psychology group at the Carl von Ossietzky University Oldenburg for their support during data acquisition.

Kirsten Weber was supported for this work by a fellowship from the Hanse Institute for Advanced Studies in Delmenhorst, Germany. The research was supported by the German Research Foundation (SFB-TRR31). The authors have no conflicts of interest to declare.

## References

Allen, K., Pereira, F., Botvinick, M., & Goldberg, A. E. (2012). Distinguishing grammatical constructions with fMRI pattern analysis. Brain and Language, 123(3), 174–182. https://doi.org/10.1016/j.bandl.2012.08.005

Arai, M., & Keller, F. (2013). The use of verb-specific information for prediction in sentence processing. Language and Cognitive Processes, 28, 525–560. https://doi.org/10.1080/01690965.2012.658072

Behrens, T. E. J., Woolrich, M. W., Walton, M. E., & Rushworth, M. F. S. (2007). Learning the value of information in an uncertain world. Nature Neuroscience, 10, 1214–1221. https://doi.org/Doi10.1038/Nn1954

Bernolet, S., & Hartsuiker, R. J. (2010). Does verb bias modulate syntactic priming? Cognition, 114, 455–461. https://doi.org/S0010-0277(09)00285-6[pii]10.1016/j.cognition.2009.11.005

Bonhage, C., Mueller, J., Friederici, A., & Fiebach, C. (2015). Combined eye tracking and fMRI reveals neural basis of linguistic predictions during sentence comprehension. Cortex, 68, 33–47. https://doi.org/10.1016/j.cortex.2015.04.011

Botvinick, M. M., Cohen, J. D., & Carter, C. S. (2004). Conflict monitoring and anterior cingulate cortex: an update. Trends in Cognitive Sciences, 8, 539–546. https://doi.org/10.1016/j.tics.2004.10.003

Boudewyn, M. A., Long, D. L., & Swaab, T. Y. (2015). Graded expectations: Predictive processing and the adjustment of expectations during spoken language comprehension. Cognitive, Affective, & Behavioral Neuroscience, 15(3), 607–624.

Brothers, T., Swaab, T. Y., & Traxler, M. J. (2015). Effects of prediction and contextual support on lexical processing: prediction takes precedence. Cognition, 136, 135–149. https://doi.org/10.1016/j.cognition.2014.10.017

Brysbaert, M., Buchmeier, M., Conrad, M., Jacobs, A.M., Bolte, J., & Bohl, M. (2011). The Word Frequency Effect. Experimental Psychology, 58, 412–424. https://doi.org/10.1027/1618-3169/a000123

Catani, M., Jones, D. K., & Ffytche, D. H. (2005). Perisylvian language networks of the human brain. Annals of Neurology, 57, 8–16.

Christiansen, M. H., & Chater, N. (2016). The Now-or-Never bottleneck: A fundamental constraint on language. Behavioral and Brain Sciences, 39, e62.

Fine, A., Jaeger, T. F., Farmer, T. A., & Qian, T. (2013). Rapid Expectation Adaptation during Syntactic Comprehension. PLoS One, 8, e77661. https://doi.org/10.1371/journal.pone.0077661

Friederici, A. D. (2009). Pathways to language: fiber tracts in the human brain. Trends in Cognitive Sciences, 13, 175–181. https://doi.org/10.1016/j.tics.2009.01.001

Friederici, A. D., & Gierhan, S. M. (2013). The language network. Current Opinion in Neurobiology, 23, 250–254.

Garnsey, S. M., Pearlmutter, N. J., Myers, E., & Lotocky, M. A. (1997). The contributions of verb bias and plausibility to the comprehension of temporarily ambiguous sentences. Journal of Memory and Language, 37(1), 58–93.

Hagoort, P. (2014). Nodes and networks in the neural architecture for language: Broca’s region and beyond. Current Opinion in Neurobiology, 28, 136–141. http://dx.doi.org/10.1016/j.conb.2014.07.013

Hagoort, P., & Indefrey, P. (2014). The Neurobiology of Language Beyond Single Words. Annual Review of Neuroscience, Vol 33, 37, 347–362. https://doi.org/doi:10.1146/annurev-neuro-071013-013847

Henderson, J. M., Choi, W., Lowder, M. W., & Ferreira, F. (2016). Language structure in the brain: A fixation-related fMRI study of syntactic surprisal in reading. NeuroImage, 132, 293–300. https://doi.org/10.1016/j.neuroimage.2016.02.050

Holdgraf, C. R., de Heer, W., Pasley, B., Rieger, J., Crone, N., Lin, J. J., … Theunissen, F. E. (2016). Rapid tuning shifts in human auditory cortex enhance speech intelligibility. Nature Communications, 7, 13654.

Kuperberg, G., & Jaeger, T. F. (2016). What do we mean by prediction in language comprehension? Language, Cognition and Neuroscience, 31, 32–59. https://doi.org/10.1080/23273798.2015.1102299

Lau, E. F., Holcomb, P. J., & Kuperberg, G. R. (2013). Dissociating N400 effects of prediction from association in single word contexts. Journal of Cognitive Neuroscience, 25(3), 484–502.

Lau, E. F., Weber, K., Gramfort, A., Hämäläinen, M. S., & Kuperberg, G. R. (2016). Spatiotemporal Signatures of Lexical–Semantic Prediction. Cerebral Cortex, 26, 1377–1387. https://doi.org/10.1093/cercor/bhu219

Loebell, H., & Bock, K. (2003). Structural priming across languages. Linguistics, 41, 791–824.

McLaren, D. G., Ries, M. L., Xu, G., & Johnson, S. C. (2012). A generalized form of context-dependent psychophysiological interactions (gPPI): A comparison to standard approaches. NeuroImage, 61, 1277–1286. http://dx.doi.org/10.1016/j.neuroimage.2012.03.068

Melinger, A., & Dobel, C. (2005). Lexically-driven syntactic priming. Cognition, 98, B11–B20. https://doi.org/10.1016/j.cognition.2005.02.001

Menenti, L., Gierhan, S. M. E., Segaert, K., & Hagoort, P. (2011). Shared Language: Overlap and Segregation of the Neuronal Infrastructure for Speaking and Listening Revealed by Functional MRI. Psychological Science, 22, 1173–1182. https://doi.org/10.1177/0956797611418347

Neely, J. H. (1991). Semantic priming effects in visual word recognition: A selective review of current findings and theories. In D. Besner (Ed.), Basic Processes in Reading and Visual Word Recognition (pp. 264–333). Hillsdale, NJ: Erlbaum.

Oldfield, R. C. (1971). The assessment and analysis of handedness: The Edinburgh inventory. Neuropsychologia, 9, 97–113.

Pickering, M. J., & Garrod, S. (2007). Do people use language production to make predictions during comprehension? Trends in Cognitive Science, 11, 105–110. https://doi.org/10.1016/j.tics.2006.12.002

Rodd, J. M., Vitello, S., Woollams, A. M., & Adank, P. (2015). Localising semantic and syntactic processing in spoken and written language comprehension: An Activation Likelihood Estimation meta-analysis. Brain and Language, 141, 89–102. https://doi.org/10.1016/j.bandl.2014.11.012

Schoot, L., Menenti, L., Hagoort, P., & Segaert, K. (2014). A little more conversation –the influence of communicative context on syntactic priming in brain and behavior. Frontiers in Psychology, 5, 208. https://doi.org/10.3389/fpsyg.2014.00208

Segaert, K., Kempen, G., Petersson, K. M., & Hagoort, P. (2013). Syntactic priming and the lexical boost effect during sentence production and sentence comprehension: An fMRI study. Brain and Language, 124, 174–183.

Segaert, K., Menenti, L., Weber, K., & Hagoort, P. (2011). A Paradox of Syntactic Priming: Why Response Tendencies Show Priming for Passives, and Response Latencies Show Priming for Actives. PLoS One, 6, e24209. https://doi.org/10.1371/journal.pone.0024209

Segaert, K., Menenti, L., Weber, K., Petersson, K. M., & Hagoort, P. (2012). Shared Syntax in Language Production and Language Comprehension—An fMRI Study. Cerebral Cortex, 22, 1662–1670. https://doi.org/10.1093/cercor/bhr249

Segaert, K., Weber, K., Cladder-Micus, M., & Hagoort, P. (2014). The influence of verb-bound syntactic preferences on the processing of syntactic structures. Journal of Experimental Psychology: Learning, Memory, and Cognition, 40, 1448–1460. https://doi.org/10.1037/a0036796

Shenhav, A., Cohen, J. D., & Botvinick, M. M. (2016). Dorsal anterior cingulate cortex and the value of control. Nature Neuroscience, 19, 1286–1291.

Snijders, T. M., Vosse, T., Kempen, G., Van Berkum, J. J. A., Petersson, K. M., & Hagoort, P. (2009). Retrieval and Unification of Syntactic Structure in Sentence Comprehension: an fMRI Study Using Word-Category Ambiguity. Cerebral Cortex, 19, 1493–1503. https://doi.org/10.1093/cercor/bhn187

Utevsky, A. V., Smith, D. V., & Huettel, S. A. (2014). Precuneus Is a Functional Core of the Default-Mode Network. The Journal of Neuroscience, 34, 932–940. https://doi.org/10.1523/jneurosci.4227-13.2014

Vassena, E., Holroyd, C. B., & Alexander, W. H. (2017). Computational Models of Anterior Cingulate Cortex: At the Crossroads between Prediction and Effort. Frontiers in Neuroscience, 11. https://doi.org/10.3389/fnins.2017.00316

Weber, K., & Indefrey, P. (2009). Syntactic priming in German–English bilinguals during sentence comprehension. NeuroImage, 46, 1164–1172.

Weber, K., Lau, E. F., Stillerman, B., & Kuperberg, G. R. (2016). The Yin and the Yang of Prediction: An fMRI Study of Semantic Predictive Processing. PLoS One, 11, e0148637. https://doi.org/10.1371/journal.pone.0148637

Wells, J. B., Christiansen, M. H., Race, D. S., Acheson, D. J., & MacDonald, M. C. (2009). Experience and sentence processing: Statistical learning and relative clause comprehension. Cognitive Psychology, 58, 250–271. http://dx.doi.org/10.1016/j.cogpsych.2008.08.002

Willems, R. M., Frank, S. L., Nijhof, A. D., Hagoort, P., & Van den Bosch, A. (2015). Prediction during natural language comprehension. Cerebral Cortex, 26(6), 2506–2516.

